# The algal PETC-Pro171-Leu suppresses electron transfer in the cytochrome *b_6_f* complex under acidic lumenal condition

**DOI:** 10.1101/2021.12.15.472847

**Authors:** Shin-Ichiro Ozawa, Felix Buchert, Ruby Reuys, Michael Hippler, Yuichiro Takahashi

**Affiliations:** Institute of Plant Science and Resources, Okayama University, Kurashiki 710-0046, Japan; Institute of Plant Biology and Biotechnology, University of Münster, 48143 Münster, Germany; Research Institute for Interdisciplinary Science, Okayama University, 700-8530, Japan

**Keywords:** photosynthesis, Q-cycle, cytochrome *b*_6_*f*, photosynthetic control

## Abstract

Linear photosynthetic electron flow (LEF) produces NADPH and generates a proton electrochemical potential gradient across the thylakoid membrane used to synthesize ATP, both of which are required for CO_2_ fixation. As cellular demand for ATP and NADPH are variable, cyclic electron flow (CEF) between PSI and cytochrome *b*_6_*f* complex (*b*_6_*f*) produces extra ATP. The *b*_6_*f* regulates LEF and CEF via photosynthetic control, which is a pH-dependent *b*_6_*f* slowdown of plastoquinol oxidation at the lumenal site. This protection mechanism is triggered at more alkaline lumen pH in the *pgr1* mutant of the vascular plant *Arabidopsis thaliana*, carrying Pro194Leu in the *b*_6_*f* Rieske Iron-sulfur protein. In this work, we introduced *pgr1* mutation in the green alga *Chlamydomonas reinhardtii* (*PETC-P171L*). Consistent with *pgr1* phenotype, *PETC-P171L* displayed an impaired NPQ induction along with slower photoautotrophic growth under high light conditions. Our data provides evidence that the ΔpH component in *PETC-P171L* is dependent on oxygen availability. Only under low oxygen conditions the ΔpH component was sufficient to trigger a phenotype in algal *PETC-P171L* where the mutant *b*_6_*f* was more restricted to oxidize the PQ pool and showed a diminished electron flow through the *b*_6_*f* complex.

**One sentence summary:** Change of PETC to P171L via site directed mutagenesis alters the pH dependency of the photosynthetic control mechanism

## Introduction

Oxygenic photosynthesis fixes carbon dioxide and generates oxygen as a result of intra- and intermolecular photochemical electron transfer reactions. The electron transfer pathways in oxygenic photosynthesis either follows Linear Electron Flow (LEF) or, as an example of alternative passages, Cyclic Electron Flow (CEF). Electrons stemming from water oxidation reduce NADP^+^ during LEF, thus providing the reducing power for CO_2_ fixation in the Calvin Benson cycle. In the CEF, electrons from reduced ferredoxin (Fd) are differently partitioned and redirected into the plastoquinone (PQ) pool. A possible CEF route reinjects electrons into the cytochrome *b*_6_*f* complex (*b*_6_*f*) from the stromal side. In LEF and CEF, the *b*_6_*f* contributes to proton gradient formation across thylakoid membrane by translocating protons from chloroplast stroma to thylakoid lumen. Light-driven charge separation, water splitting as well as *b*_6_*f* activity generate a membrane potential (ΔΨ) and a proton gradient (ΔpH), respectively, and this proton motive force (*pmf*) drives ATP synthesis. During prolonged electron transfer, the lumen will get further acidified, which in turn will slow down plastoquinol (PQH2) oxidation at the Qo-site of *b*_6_*f*, and thereby establish photosynthetic control to protect photosystem I (PSI) from acceptor side limitation (Rumberg and Siggel, 1969; Tikhonov et al., 1981; Nishio and Whitmarsh, 1993). Lumen acidification is also important to induce Non-Photochemical Quenching (NPQ) (Kanazawa and Kramer, 2002), which is the most rapid photoprotective response to excess light. The fastest constituent of NPQ is qE, which operates at a time scale of seconds to minutes and regulates the thermal dissipation of excess absorbed light energy, thereby providing effective photo-protection of photosystem II (PSII). In vascular plants, the PSII protein PSBS is essential for qE (Li et al., 2002) whereas qE induction in the green alga *Chlamydomonas reinhardtii* is mediated by LHCSR3, an ancient light-harvesting protein that is missing in vascular plants (Peers et al., 2009).

Electron transfer through the *b*_6_*f* is governed by the light-induced introduction of an electron hole (positive charge) into the high-potential chain. The latter involves redox centers heme *f* and the [2Fe-2S] type Rieske Iron Sulfur Cluster (Rieske ISC). The electron hole depends on photo-oxidized plastocyanin and is successively passed from heme *f* via Rieske ISC to PQH_2_. The Q-cycle within the *b*_6_*f* is responsible to generate a *pmf* across the thylakoid membrane, owing to a sophisticated electron bifurcation. Two electrons are pulled from PQH_2_ at the Q_o_ site, forming PQ within the membrane while simultaneously releasing two H^+^ into the lumen. During the PQH_2_ oxidation at the Q_o_ site, the first electron is transferred into the high-potential chain (Rieske ISC reducing heme *f*), and the second electron into the low-potential chain (*b*_L_ heme reducing *b*_H_ heme, terminal acceptor is *C*_i_ heme). A second PQH_2_ oxidation at the Q_o_ site forms a reduced form of *b*_H_ / *C*_i_ heme pair which reduces a PQ bound at Q_i_ site to complete the Q-cycle (note that alternative annotations for Q_o_, Q_i_, *b*_H_ and *C*_i_ can be found in literatures). It was suggested that under LEF and CEF conditions two types of Q-cycle mechanism operate in the *b*_6_*f* with specific lumen acidification efficiencies per electron transferred via the high-potential chain (Buchert et al., 2020).

The molecular structure of *b*_6_*f* complex was first resolved in the green alga *Chlamydomonas reinhardtii* and the cyanobacteria *Mastigocladus laminosus* independently (Kurisu et al., 2003; Stroebel et al., 2003). The *b*_6_*f* functions as a homodimer, and each monomer consists of four core subunits (PetA, PetB, PETC, and PetD) and four stably associated small subunits (PetG, PetL, PETM, and PETN) to stabilize the complex (Figure 1A). The cofactors and quinone binding pockets involved in electron transfer are found in the *b*_6_*f* core subunits which resemble the cytochrome *bc*_1_ complex of the respiratory chain. The structure-function similarities suggest that the hydrophilic domain of PETC subunit would move toward PetA subunit in transferring the electrons from Rieske ISC to heme *f*.

**Figure 1.**
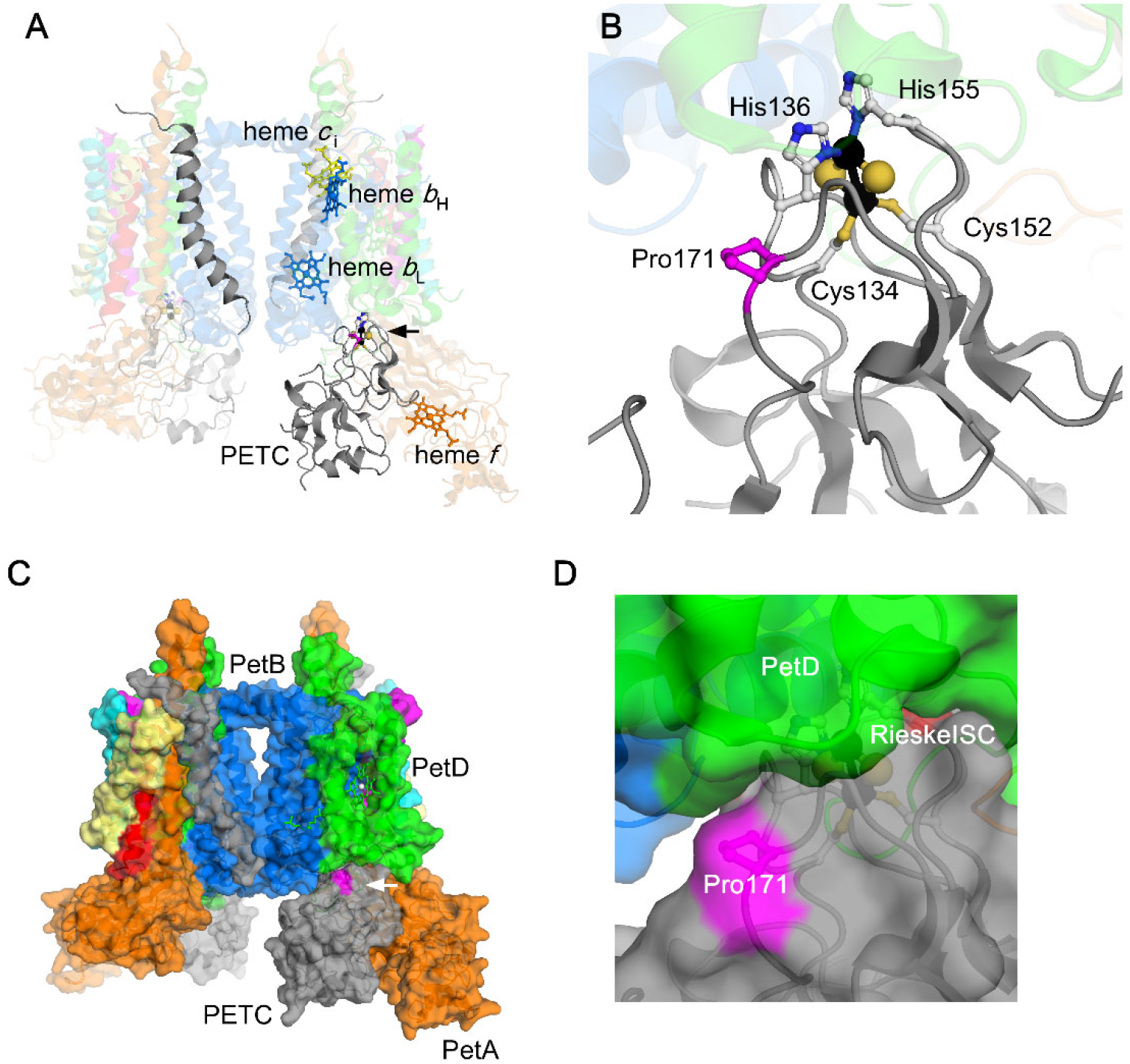
The position of Rieske ISC and PETC-Pro171 in the Chlamydomonas *b*_6_*f* complex. Structures of the dimeric *b*_6_*f* from the *C. reinhardtii* (PDB; 1Q90). The flexible hinge of PETC is not modeled. The Rieske ISCs are shown in space filling representation and of which ligand amino acids (Cys134, His136, Cys152, and His155 of PETC) and Pro171 of PETC (colored in magenta) and four hemes (heme *c*_i_ is yellow, heme *b*_H_ and heme *b*_L_ are blue, and heme *f* is orange) are shown in ball-stick model. The PetA (orange), PetB (blue), PETC (gray), PetB (green), PetG (magenta), PetL (yellow), PETM (cyan), and PETN (red) subunits are shown. **(A)** and **(C)**. Overall structure with the lumenal side at bottom and the stromal side at the top. The position of one of the Rieske ISC is indicated with arrow. **(B)** and **(D)**. The enlarged view around the Rieske iron sulfur cluster. **(C)** and **(D)**. Protein surface is modeled. The subunits and cofactors except for PETC subunit are transparent in panel A and B.

In *Arabidopsis thaliana*, various photosynthetic mutants have been isolated and characterized by forward genetics. One study employed ethyl methanesulfonate as mutagen to isolate the CE10-10-1 mutant which exhibited reduced NPQ with normal PSI/PSII quantum yields (Shikanai et al., 1999). A follow-up study pinpointed the mutation as a proline-to-leucine substitution at position 194 of Arabidopsis PETC, linking the NPQ defect to insufficient ΔpH formation in the light (Munekage et al., 2001). The Arabidopsis PETC-P194L mutation was dubbed *pgr1* (*<u>p</u>roton <u>g</u>radient <u>r</u>egulation* 1).

It is known that green algal photosystems are protected from excess light illumination differently than plants. For instance, algal state transitions redistribute excitation energy more effectively than in plants (Delepelaire and Wollman, 1985). Moreover, flavodiiron protein was found in algae to protect PSI acceptor side limitation induced by fluctuating light illumination (Jokel et al., 2018). Induction of qE relies on LHCSR3, which requires a preceding stress period to trigger light to heat dissipation upon lumen acidification (Peers et al., 2009; Bonente et al., 2011). Moreover, the algal PETO protein interacts with *b*_6_*f* and functions as a CEF effector under anoxic condition, which might point to a different CEF regulation (Takahashi et al., 2016; Buchert et al., 2018).

Herein, we introduced the Arabidopsis *pgr1* mutation into *Chlamydomonas reinhardtii* (*PETC-P171L*) to analyze the effects of the mutation on Photosynthetic Control in the green alga. The *PETC-P171L* cytochrome *b*_6_*f* showed a normal accumulation and electron flow activity in cells grown under oxic conditions. However, the *PETC-P171L* displayed an impaired photoautotrophic growth and was less competent in NPQ induction. Unlike Arabidopsis *pgr1*, the *PETC-P171L* phenotype was conditional and revealed upon cellular stress treatment. In line, the electron transfer activity of the mutated *b*_6_*f* was significantly decreased in cells grown under anoxic conditions. Our experiments also revealed that the ΔpH component in *PETC-P171L* is dependent on oxygen availability under light conditions that are permissive under other circumstances. Under low oxygen condition the mutant *b*_6_*f* was more restricted to oxidize the PQ pool and limited electron flow.

## Materials and methods

### Strains and growth condition

The cells of *Chlamydomonas reinhardtii* were grown in Tris-acetate phosphate (TAP), Tris-Phosphate (TP) medium, or minimum medium (MM) (Harris, 1988) under 50-100 μmol photons·m^−2^·s^−1^ or 600 μmol photons·m^−2^·s^−1^. The previously isolated PETC gene disrupted strain *petC-Δ1* (mating type minus) described in (de Vitry et al., 1999) was crossed with cc-125 (mating type plus) and the resultant PETC deficient progeny was used as a recipient strain for transformation. Doubling time of the cell growth is calculated by dividing 0.30 by specific growth rate obtained as the slope which is computed by linear approximation for logarithmic growth curve in common logarithmic axis. Linear fitting was performed at the highest coefficient of determination (R-square) with no restrictions by using built-in linear fitting program of the Origin Pro 2021b version 9.8.5.212 (OriginLab corporation, USA). Refer the Appendix 1 for derivation of the formula to calculate the doubling time with specific growth rate. For *in vivo* spectrophotometric analysis, cells were maintained at 20 μmol photons·m^−2^·s^−1^ on agar-supplemented TAP plates (Sueoka, 1960). When growing cells for experiments, liquid TP medium was used. The cultures were grown at 40 μmol photons·m^−2^·s^−1^ (16 h light/8 h dark) and were bubbled with sterile air at 25°C while shaking at 120 rpm. About 300 ×10^6^ cells were incubated in 400-mL TP in a FMT 150 photobioreactor vessel (Photon Systems Instruments, Czech Republic). Photobioreactor experiments were carried out in the absence of air supply and comprised 3 days of 16-h photoperiods (sigmoidal light and temperature gradients of 0-250 μmol photons·m^−2^·s^−1^ from white-red LEDs and 18-28°C, respectively, with 18°C constant in the dark).

### Generation of transformation vectors

The BamHI fragment containing *PETC* gene (Cre11.g467689.t1.1, Chlamydomonas version 5.5) in the pACR4.5 (de Vitry et al., 1999) was digested with BamHI and subsequently the resulting 4153 bp fragment containing the *PETC* gene and 528 bp upstream of 5′ -UTR of *PETC* was recloned into pBSSK^−^vector at BamHI site to yield pgcC plasmid. The mutation Pro171Leu in PETC was introduced by In-Fusion^R^ (Clontech, TAKARA, Japan) with the DNA fragment between BbvCI and SnaBI sites which was generated by PCR with If-BbvCI_P171L_gcC_F and If-SnaBI_gcC_R primers. The sequence of this study’s oligos is shown in Table 1. The DNA sequence of the inserted PCR product was validated by DNA sequencing with the primers used to generate the PCR product and we obtained pgcC-P171L plasmid.

We also prepared the DNA construct in which the *PETC* gene expression is controlled by Chlamydomonas *PSAD* promoter and UTRs. We amplified spliced *PETC* coding DNA sequence fragment by nested PCR with cDNA library as a template DNA. The 1st round of PCR from cDNA library with primers HK#1483 and HK#1484 generated a 731 bp DNA fragment. The following 2nd round of PCR from the 1000 times diluted 1st round of PCR product with primers HK#1485 and HK#1486. The resulting DNA fragment of the PCR was ligated into the pSL18 vector (Depège et al., 2003) digested with NdeI and XbaI. The cloned DNA sequence of spliced *PETC* coding DNA sequence was validated by DNA sequencing with HK#1491 and HK#1492 primers. To introduce Pro171Leu in PETC, the *PETC* coding DNA sequence was replaced by In-Fusion HD cloning kit at NdeI and XbaI site with the PCR product consisting of two overlapping DNA fragments generated by the following primers; one fragment is generated with If-paDcC-NdeI_F and ccC_mP171L_R while the other fragment is generated with ccC_mP171L_F and If-paDcC-XbaI_R. The replaced DNA sequence was validated by DNA sequencing with the primers used to generate PCR product and cC_cdn131-137_F and cC_cdn137-131_R, and we obtained pccC-P171L plasmid.

### Nuclear gene transformation

Nuclear gene transformation was performed with NEPA21 (Nepagene, Japan) according to the previous research (Yamano et al., 2013). The plasmid DNA was linearized by digesting with ScaI and the digested DNA was delivered into the *ΔPETC*. Selection of transformants was performed on MM solid medium under continuous illumination (50-100 μmol photons·m^−2^·s^−1^).

### SDS-PAGE

The whole cellular proteins were denatured by heating at 100°C for 2 minutes in the presence of SDS and DTT and the polypeptides were separated by SDS-PAGE. The polypeptides were transferred on nitrocellulose membrane for immunodetection. The signals were visualized with a CCD camera equipped imagers (Amersham Imager, LAS-4000, GE healthcare, or Chemidoc, Biorad) and the signal intensities were quantified with Multigauge ver. 3.0 (FujiFilm, Japan)

### Low temperature fluorescence emission spectra

Cells were resuspended in TP medium at 2 μg Chl·mL^−1^ and incubated for overnight under illumination at 300 μmol photons·m^−2^·s^−1^ prior to measurements. After the incubation, culture was directly, state 1 inducible treatment, or state 2 inducible treatment at 25°C. For the case of state 1 inducible treatment, cells were incubated under 300 μmol photons·m^−2^·s^−1^ in the presence of 10 μM of 3-(3,4-dichlorophenyl)-1,1-dimethylurea (DCMU). For the case of state 2 inducible treatment, cells were incubated in the darkness in the presence of 100 mM glucose and 50 U·mL^−1^ of glucose oxidase. A sample in a NMR tube was in liquid nitrogen, excited with 430 nm illumination, and emission spectrum was recorded from 400 to 800 nm with FL-7000 (Hitachi, Japan).

### *In vivo* Spectroscopy

TP-grown cells with sterile air supply were harvested after ~2-h in the light cycle. Grown cultures were diluted ~6-fold at least once after inoculation and grown to a density of ~1×10^6^ cells·mL^−1^ before harvesting (5000 rpm, 5 min, 25°C). Cells were resuspended at 20 μg Chlorophyll·mL^−1^ in TP supplemented with 20% (*w*/*v*) Ficoll. Oxic samples were resuspended throughout the measurements in 2 mL open cuvettes. For oxygen-deprived conditions, cells were supplemented with 50 mM glucose, 10 U glucose oxidase (Carl Roth, Germany) and 30 U catalase (Sigma-Aldrich, USA) in the cuvette, and then overlaid with mineral oil for 20 min in the dark. The enzyme stock solutions to obtain anoxic samples were freshly prepared. Cell incubation in the presence of 10 μM nigericin (from 10 mM ethanolic stock) was carried out for 20 min in the light before the measurements. All samples were light-adapted in the cuvette for 20 min before the measurements using LEDs emitting ~150 μmol photons·m^−2^·s^−1^ of 630 nm actinic light (AL). Measurements of the electrochromic shift (ECS) and P700 were obtained using a Joliot-type spectrophotometer (JTS-10, Biologic France in Figure 3; otherwise JTS-150, Spectrologix USA) and are described in detail elsewhere (Buchert et al., 2020; Buchert et al., 2022). In brief, all absorption changes were normalized to the ECS ΔI/I signals (520 nm – 546 nm) produced after a saturating (6-ns laser flash in the presence of 1 mM hydroxylamine (HA) and 10 μM DCMU, which relate to the density of active PSI centers in the sample. Thus, the unit of 1 charge separation·PSI^−1^ is produced when measuring the flash-induced ECS amplitude (rapidly established before the first detection at ~300-μs). This optical ruler was determined in separate measurements to compare sample activities regardless of the presence of HA and DCMU. The shown flash-induced ECS kinetics were obtained after lowering the flash intensity to 30-45% saturation to avoid double turnovers of the cytochrome *b*_6_*f* complex. The flash-induced ECS kinetics are multiphasic and the *b*-phase, which is a ~10-ms ECS rise associated with charge separation activity via the *b*_6_*f*, was deconvoluted to obtain the shown half-times. Deconvolution from the raw ECS kinetics involved subtraction of rate constants of a two-exponential decay function to fit the *c*-phase, which is associated with ATP synthase activity. The deconvoluted *b*-phase was quantified with a mono-exponential function (Buchert et al., 2020). The ECS was also deconvoluted in the dark pulse method (Joliot and Joliot, 2002; Nawrocki et al., 2019). Accordingly, AL-adapted cells experienced 30 s darkness and were re-illuminated with AL (Supplemental Figure 1). During this 10 s AL induction phase, activities of stable charge separations·s^−1^PSI^−1^ were measured by brief light shuttering at discrete time points (four technical replicates with 1 min AL in between; data assembly from two independent measurement sequences). For AL interruptions at *t* = 0 ms, the linear ECS slopes before (S_L_ from −4 ms to −1 ms) and after darkness (S_D_ from +1 ms to +5 ms) gave electron transfer rates before darkness (S_L_ – S_D_). Rates at 1 ms illumination of light-adapted cells were calculated from separate slopes obtained during 2 ms AL after 30 s dark (15 replicates spaced by 5 s). P700 measurements (705 nm – 740 nm) are based on the method by (Klughammer and Schreiber, 1994), measuring AL-adapted cells. As shown in Supplemental Figure 2, the fractions of photooxidizable (by a 22 ms saturating pulse), pre-oxidized in AL and non-photooxidizable P700 were determined by referring to the fully oxidized P700 signals in PSII-inhibited HA/DCMU samples (Buchert et al., 2020; Buchert et al., 2022). Since the above-mentioned ECS slopes obtained during 2 ms AL are a function of the P700 oxidation rate in PSII-inhibited HA/DCMU samples, the obtained YI parameter in those samples served to calculate CEF (by multiplication with the P700 oxidation rate) as it a function of the concentration of reduced P700 in AL (Takahashi et al., 2013). Single-turnover measurements of the cytochrome *b*_6_*f* complex were measured after a sub-saturating flash, hitting 40-50% of PSI reaction centers (Buchert et al., 2020; Buchert et al., 2022). Using interference filters, the kinetics of flash-induced cytochrome-*f* reduction (measured with 554-nm detection pulses) was calculated using a baseline drawn between 546-nm and 573-nm after correcting for the spectral overlap with the *b*-hemes by subtracting 0.27 × (563-nm – 546-nm).

### Chlorophyll fluorescence measurement

Cells were grown in liquid TP medium to ~1 × 10^6^ cells·mL^−1^ with gentle shaking at 25°C under continuous light (50 μmol photons·m^−2^·s^−1^). The cells were harvested by centrifugation at 4000 rpm, suspended in fresh liquid TP medium at 5 μg Chl·mL^−1^, and placed in darkness for 30 min (growth light control samples). Photosynthesis driven by 450 nm LEDs was measured with a Maxi-Imaging PAM chlorophyll fluorometer (Heinz Walz, Germany). The effective photosynthetic active radiation in the measuring chamber was determined with a Quantitherm PAR/Temp Sensor (Hansatech, England). Cell aliquots were also incubated shaking for 4 h under high light illumination (500 μmol photons·m^−2^·s^−1^). The maximal PSII quantum yield in dark-adapted samples and the PSII quantum yield during illumination Y(II) were determined as (Fm–Fo)/Fm and (Fm’–F)/Fm’, respectively. Fo and Fm of dark-adapted samples are respective minimal (in the presence of detection light) and maximal fluorescence yields (due to closed PSII centers after saturating pulse). F and Fm’ are the fluorescence yields in light-adapted cells obtained before and after a saturating pulse, respectively. The nonphotochemical quenching NPQ = (Fm–Fm’)/Fm’ was also determined, as well as estimation of the plastoquinone pool redox state using the (1–qL) parameter (Kramer et al., 2004). The latter required Fo’ measurements (minimal fluorescence in illuminated samples) and qL was calculated as ((Fm’–F)×Fo’)/((Fm’–Fo’)×F). These parameters were either measured after 30 s during light curve experiments or at distinct time points when a 10 min continuous illumination at 95 μmol photons·m^−2^·s^−1^ was applied. Fluorescence data in FMT 150 photobioreactor experiments was obtained from blue excitation lights, and Y(II) was calculated as above using saturating pulses every 15 min. The NPQ values were obtained from calculations with OD_680_-corrected Fm and Fm’ whereas for each 16-h photoperiod the respective Fm was averaged over a 2-h dark period before the onset of light.

## Results

The PETC protein consists of one transmembrane helix and a lumenal hydrophilic domain in which the Rieske iron sulfur cluster (ISC) is present, and both domains are connected through a flexible hinge. Based on the structural similarities with the Rieske subunit in *bc*_1_ complex of respiratory chain, photosynthetic electron transfer between Rieske ISC and heme *f* in PetA involves reversible movements of the lumenal hydrophilic domain of PETC. The PETC amino acid sequence alignment shows that chloroplast targeting transit peptide of *A. thaliana* is 23 amino acids longer than that of *C. reinhardtii* while the mature protein sequence is highly conserved (Supplemental Figure 3). The four amino acids for ligation of two iron atoms and a flexible hinge region are conserved. Moreover, the mutated proline residue in Arabidopsis *pgr1* (Pro194) is conserved as Pro171 in *C*. *reinhardtii*. The location of this proline is close to Rieske ISC and is on the surface of hydrophilic domain (Figure 1B, C, and D), and its proximity to the Rieske ISC is conserved in the *b*_6_*f* structures (Supplemental Figure 4).

### Generation of *PETC-P171L* mutant in the green alga *Chlamydomonas reinhardtii*

The Arabidopsis *pgr1* mutant carrying a single amino acid alteration mutation in PETC (P194L) exhibits a hypersensitive photosynthetic control and downregulates the electron transfer flow under strong light illumination more strongly than wild type (Munekage et al., 2001). To characterize the corresponding PETC mutation in the green alga *C. reinhardtii*, we generated a *PETC-P171L* strain in which proline 171 is substituted by leucine. In our complementation-based screening approach, *petC-Δ1* was selected as a recipient strain since it has a frameshift upon TG base deletions within the 36^th^ codon and does not accumulate *PETC* transcripts, thus showing no spontaneous reversion (de Vitry et al., 1999). Since the *petC-Δ1* exhibited a reduced accumulation of PSI protein, we backcrossed the *petC-Δ1* to wild type strain (cc-125) to assess whether the mutation of PETC gene affects the accumulation of PSI proteins. We obtained several tetrads and analyzed the accumulation of PsaA and PETC by immunoblotting (Supplemental Figure 5). The T_15-3_ progeny accumulates no PETC but PsaA normally and is referred to as the ΔPETC for the subsequent transformation since the two phenotypes segregated. As described in Materials and Methods, we generated two types of vectors for transforming *PETC* gene; one consists of genomic DNA fragment around PETC gene and the other contains PETC cDNA fused with *PSAD* promoter and UTRs. We delivered each transforming vector into the ΔPETC cells by electroporation and obtained numerous colonies after screening under photoautotrophic condition (on minimum solid medium in the light of 50 μmol photons·m^−2^·s^−1^). The resulting transformants recovered the accumulation of PETC to wild-type level, indicating that both vectors efficiently complemented the ΔPETC phenotype (Figure 2). Subsequently, we transformed ΔPETC with vectors containing the mutation PETC-P171L and obtained complemented strains expressing the mutated PETC. Immunoblot analyses revealed that the complemented PETC-P171L cells accumulated PETC as well as PetA at wild-type levels, suggesting that the mutation does not affect the stability of *b*_6_*f* complex.

**Figure 2.**
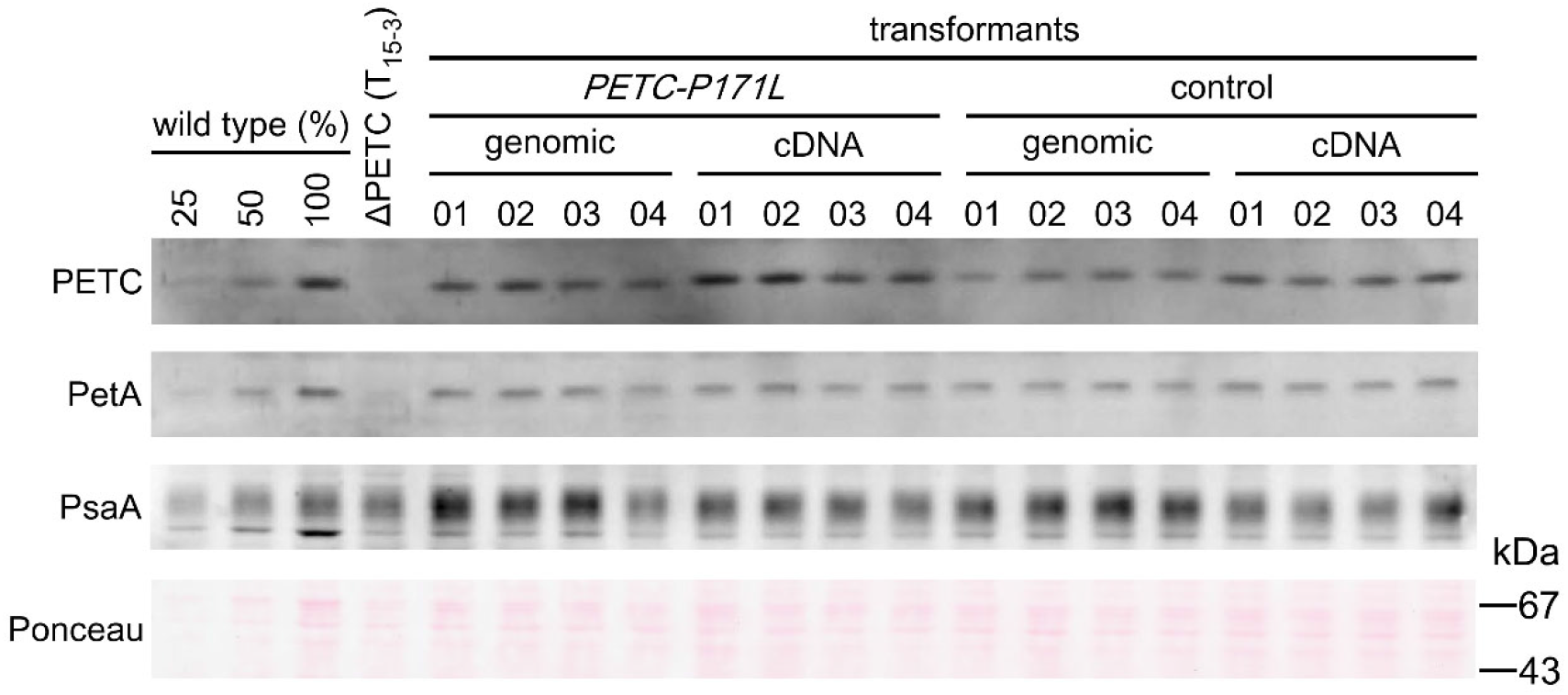
Generation of *PETC-P171L* mutant. The accumulation of PSI (PsaA) and *b*_6_*f* (PETC and PetA) proteins was analyzed by immunoblotting. The cells were grown in TAP liquid medium under the 50 μmol photons·m^−2^·s^−1^. The Pro171Leu of PETC (*PETC-P171L*) and wild type sequence of PETC (control), the PETC gene expressing with own promoter and UTRs (genomic) or the spliced PETC gene expressing with PSAD promoter UTRs (cDNA). Four independent clones from each transformant, a dilution series for the wild type (wild type (%)), and ΔPETC (T_15-3_ in Supplemental Figure 5) were also analyzed.

### The *PETC-P171L* accumulates more oxidizable P700 and displays slower onset of electron transfer in anaerobic algae which relied on the lumen pH under these conditions

To examine the *PETC-P171L* mutant *in vivo* we chose a moderate light intensity for the following experiments (~160 μmol photons·m^−2^·s^−1^ actinic red light). Moreover, *pmf* partitioning was modulated on a metabolic level in light-adapted cells by lowering oxygen availability. A similar situation may occur in the natural habitat of Chlamydomonas which shares it with respiring microbes. Accordingly, the alga faces microoxic environments in which photosynthesis needs to be fine-tuned, which is consistent with its versatile anoxic metabolism (Grossman et al., 2011). Figure 3 shows a spectroscopic *in vivo* analysis of control and the *PETC-P171L* in oxic conditions and upon anaerobic treatment. Mitochondrial congestion in anoxic samples coincides with a redox poise shown to trigger CEF (Alric, 2014) and a *pmf* re-partitioning towards a more pronounced ΔpH (Finazzi and Rappaport, 1998).

Figures 3A and 3B display the redox state of P700, the primary electron donor of PSI (see Materials and Methods and Supplemental Figure 2 for representative kinetics). Under oxic conditions when PSII was active, the P700 redox states were indistinguishable between control and *PETC-P171L* (Figure 3A). About 20% of P700 were oxidized during actinic background light which is ascribed to similar donor side limitation (YND) in both strains. P700 was able to be further photooxidized by a short saturating light pulse, as shown by the yield of PSI (YI). A similar fraction (about 40%) of P700 was not oxidizable due to acceptor side limitation (YNA). On the contrary, YND was remarkably large when PSII was inhibited (HA+DCMU, Figure 3A) because electrons from PSII were suppressed. This coincided with YI of ~0.2 in control, i.e., 20% of P700 remained reduced in the light-adapted state in absence of PSII activity and were only oxidized by saturating pulses. This finding suggests that this 20% of P700 is reduced by electrons through CEF. YI in controls was roughly twice as large as in *PETC-P171L*, which translates in lower CEF rates in the mutant (indicated as 12 electrons·s^−1^·PSI^−1^ in parenthesis of Figure 3A).

**Figure 3.**
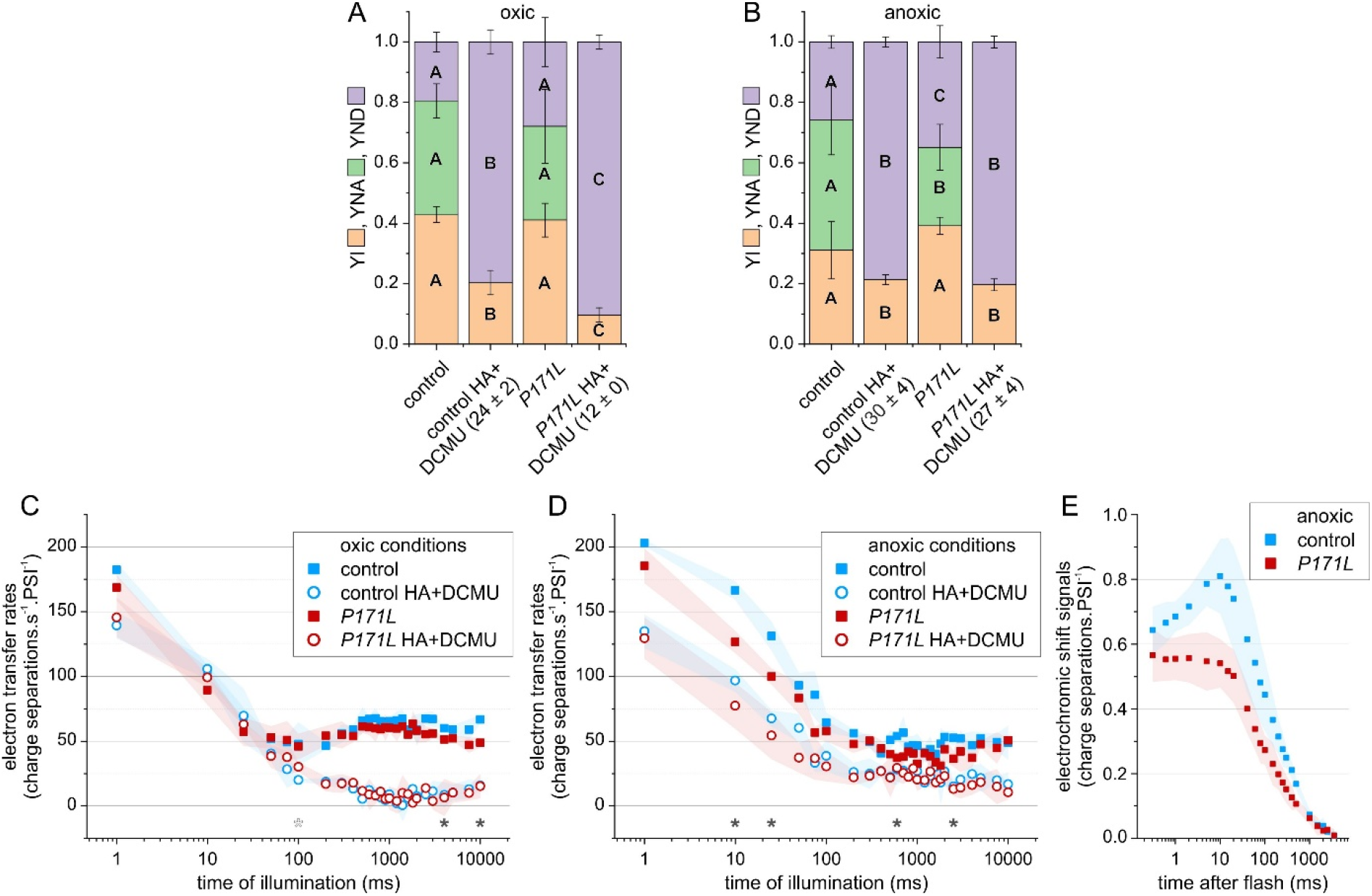
The *PETC-P171L* mutation restricts photosynthetic performance of light- and dark-adapted cells predominantly in anaerobic conditions. P700 redox states were measured in **(A)** oxic and **(B)** anoxic light-adapted cells. The fractional yields of P700 in the photo-oxidizable state (YI), the non-oxidizable form due to acceptor side limitation (YNA), and the pre-oxidized P700 due to donor side limitation (YND) are shown (parameter-specific one-way ANOVA, *n* = 3, *P* < 0.05). The oxic controls in absence of PSII activity (HA+DCMU) retained a higher YI due to enhanced steady state CEF (expressed charge separations·s^−1^·PSI^−1^ ± SD). In anoxic conditions, *PETC-P171L* failed to establish the YNA due to larger YND. **(C)** After a 30-s dark period, electron transfer rates over a 10-s photosynthetic induction phase were measured in oxic light-adapted cells. The kinetics were largely indistinguishable (means ± SD are shown, *n* = 3). Total electron transfer rates in the controls were enhanced at a later stage of illumination (filled asterisks, Student’s t-test *P* < 0.05), and *PETC-P171L* showed transiently higher rates at 100-ms in absence of PSII activity (open asterisk). **(D)** Total electron transfer in *PETC-P171L* was significantly less efficient at discrete time points, especially in the initial ~25-ms (filled asterisk). A significant difference during induction required PSII activity. **(E)** The electrochromic shift ~10-ms rising phase, associated with charge separation activity upon a single *b*_6_*f* turnover, was less pronounced in anoxic light-adapted *PETC-P171L*. Sub-saturating flashes were fired on light-adapted cells after a 30-s dark period.

In anoxic conditions the *PETC-P171L* established a larger YND and a lower YNA compared to control when PSII was active (Figure 3B). In the absence of PSII activity, control and *P171L* showed similar P700 parameters. This finding shows that CEF rates in the steady state were unaffected in both strains (shown as electrons·s^−1^·PSI^−1^ in parenthesis of Figure 3B). Overall, we observed subtle differences on the level of light-adapted steady-state of P700. The differences were produced in the absence of PSII activity under oxic conditions (Figure 3A) and in the presence of PSII activity under anoxic conditions (Figure 3B). We concluded that the PSI donor side of the *PETC-P171L* is specifically limited in anoxic condition.

Next, we re-illuminated cells after a 30-s dark period. As mentioned above, a significant ΔpH is generated when algae are kept in the dark under anoxic conditions (Finazzi and Rappaport, 1998), which supposedly slows down the Qo-site activity in *PETC-P171L*. Panels C – E in Figure 3 tested this assumption. The electrochromic shift (ECS) of photosynthetic pigments allows to measure the membrane potential as the ECS signals linearly follow the electric *pmf* component (Junge and Witt, 1968; Witt, 1979). More specifically (see Materials and Methods), we measured electron transfer transients during the first 10-s of light in oxic (Figure 3C) and anoxic conditions (Figure 3D). The induction kinetics of total electron transfer rates were similar in both strains under oxic conditions (Figure 3C). The sum of LEF and CEF activities in the control strain was slightly higher at 4-s and 10-s of light (filled asterisks, Figure 3C). The kinetics were almost identical when PSII was inhibited, except for 100-ms of light where *PETC-P171L* produced slightly higher rates (open asterisk, Figure 3C). The induction of total electron transfer under anoxic conditions showed that activities were transiently less efficient in *PETC-P171L*, producing the largest differences within the first 25 ms of light (cf. filled symbols and asterisk in Figure 3D). When PSII was inhibited (open symbols Figure 3D), no differences were observed between both strains.

We noticed that light-adapted, anoxic cells produced different ECS kinetics when a sub-saturating flash was fired after 30-s darkness (Figure 3E). The measurements resolved the averaged *b*- and *c*-phases of the flash-induced ECS kinetics (reviewed in Bailleul et al., 2010). The non-resolved *a*-phase with an amplitude of ~ 0.6 charge separations·PSI^−1^ relied on the charge separation activity of both photosystems. The *b*-phase, which is a ~10-ms ECS rise, involves charge separation activity via the *b*_6_*f* and the *c*-phase depends on ATP synthase activity. The latter phase overlapped the *b*-phase since ATP synthase activity was driven by the *pmf* provided from photosystem activity. Under oxic conditions, the fast *c*-phase prevented the appearance of a *b*-phase (Supplemental Figure 6). Therefore, the detection of an apparent *b*-phase requires a slow *c*-phase in anoxic algae (Velthuys, 1978). The ECS kinetics in anoxic *PETC-P171L* displayed a flat *b*-phase and comparably slow *c*-phase kinetics (Figure 3E), suggesting a compromised reaction of *b*_6_*f* electron transfer activity in *PETC-P171L*.

This observation prompted us to measure cytochrome-*f* reduction kinetics in single-turnover measurements. In line with the conditional phenotype of *PETC-P171L*, we did not observe a difference in oxic cells (Supplemental Figure 7). When cells were treated under anoxic conditions, we observed that the half-time of cytochrome-*f* reduction in the control strain was significantly faster than in *PETC-P171L* (43 ±23 ms vs. 74 ±8 ms, Figure 4A). To test whether this difference is linked to the ΔpH generated by anoxic algae in the dark (Finazzi and Rappaport, 1998), we took advantage of nigericin. The H^+^/K^+^ ionophore exchanges the osmotic *pmf* component (ΔpH) for the electric one (ΔΨ), and indeed it accelerated cytochrome-*f* reduction in both strains to the same level (Figure 4A). Although direct nigericin action may cause an electrogenic bias, the drug also restored the generation cytochrome *b*_6_*f*-associated charge separation phase during the first 10-ms after the flash (this so-called *b*-phase is shown as solid lines in Supplemental Figure 8). The *b*-phase half-times in *PETC-P171L* were significantly slower compared to the control strain (Figure 4B). In contrast, the *b*-phase buildup was accelerated by nigericin treatment. However, it was slightly delayed in nigericin-treated *PETC-P171L* compared to the control strain. Although nigericin also accelerated the *b*-phase buildup in the control strain, the difference was not significant.

**Figure 4.**
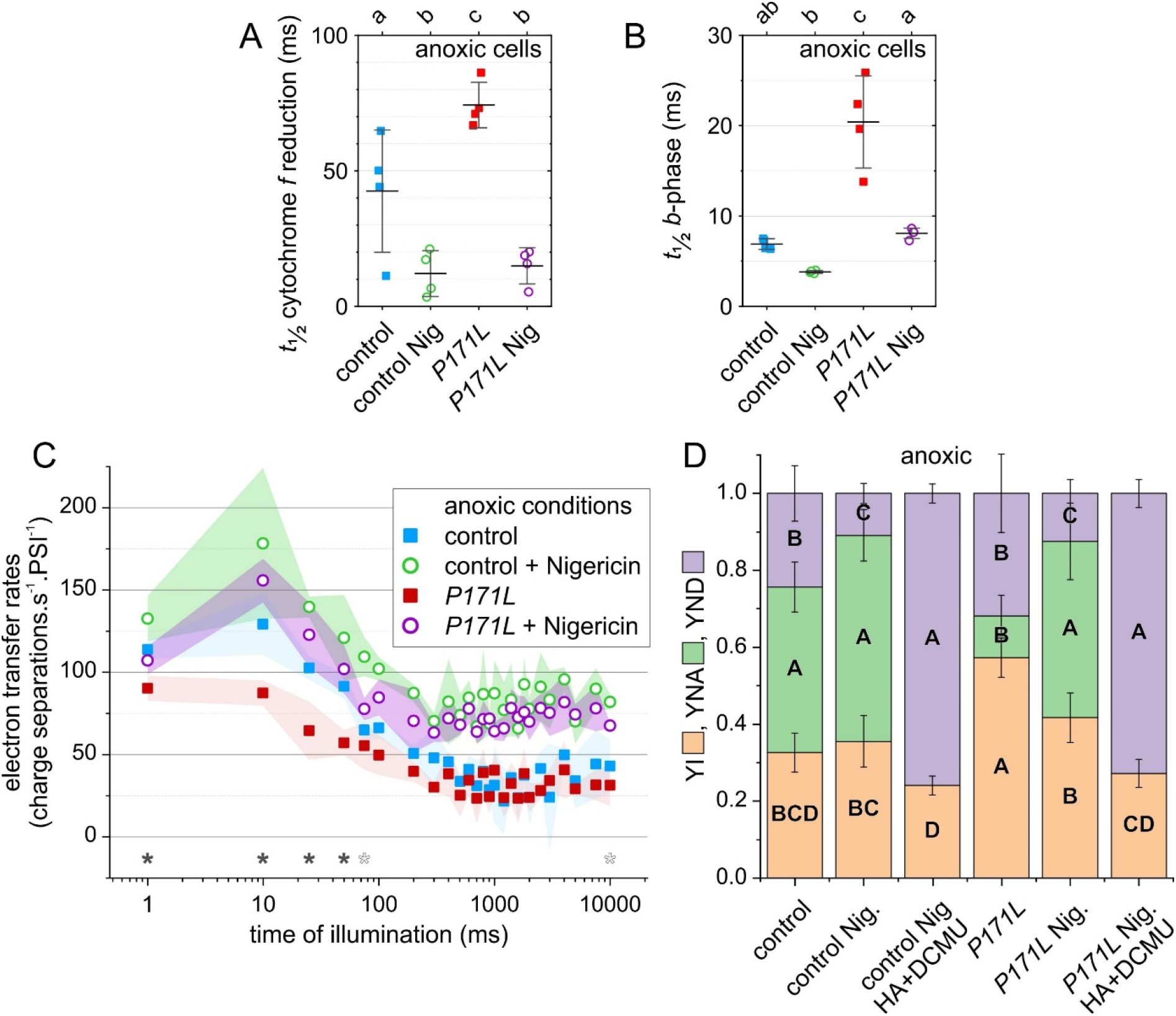
The cytochrome *b*_6_*f*-related electron transfer restrictions in *PETC-P171L* cells under anaerobic conditions is overcome by diminishing the ΔpH via nigericin. **(A)** The control and *PETC-P171L* cells displayed different cytochrome-*f* reduction half-times upon a single cytochrome *b*_6_*f* turnover under anoxic conditions (means of n=4 ±SD are shown, miniscules on top indicate significances using One-Way ANOVA/Fisher-LSD, P > 0.05). The slower cytochrome-*f* reduction in *PETC-P171L* cells as well as control strain kinetics were accelerated to similar values upon nigericin (Nig) addition. **(B)** The events that affected cytochrome-*f* reduction also decreased the *PETC-P171L* electrogenic activity in the low-potential chain during the Q-cycle. The *b*-phase (for kinetics, see Supplemental Figure 8) also showed a longer half-time in *PETC-P171L* (means of n=4 ±SD are shown, miniscules on top indicate significances using One-Way ANOVA/Fisher-LSD, P > 0.05). Nigericin addition accelerated the buildup of the *b*-phase in the *PETC-P171L* strain. The drug accelerated the *b*-phase insignificantly in the control. **(C)** After a 30-s dark period, electron transfer rates over a 10-s photosynthetic induction phase were measured in anoxic light-adapted cells (averaged kinetics ±SD, *n* = 3). Total electron transfer in *PETC-P171L* was significantly less efficient within the first ~50-ms (filled asterisk, Student’s t-test *P* < 0.05). The different induction kinetics between the strains relied on the ΔpH and became similar after addition of the H^+^/K^+^ ionophore nigericin (Nig., 10 μM), although the control showed higher rates at discrete sampling intervals (open asterisk). In both strains the steady state rates were higher when Nig. was present. **(D)** On the level of co-dependent PSI parameters, the fractional yields of P700 in the photo-oxidizable state (YI), the non-oxidizable form due to acceptor side limitation (YNA), and the pre-oxidized P700 due to donor side limitation (YND) are shown (parameter-specific one-way ANOVA, *n* = 3, *P* < 0.05). The YI was larger in *PETC-P171L* owing to a diminished YNA. YND was lowered to similar levels after adding Nig. and diminishing ΔpH-dependent photosynthetic control. Moreover, further PSII inhibition (HA+DCMU) produced similar P700 yields.

In the light of the nigericin effect on the single-turnover cytochrome-*f* kinetics and *b*-phase generation, we further examined the phenotype of anoxic *PETC-P171L* cells in multiple-turnover conditions. To do so, we repeated the electron transfer (Figure 4C) and P700 experiments (Figure 4D) in the presence and absence of nigericin. Indeed, Figure 4C shows that nigericin action abolished the *PETC-P171L* electron transfer bottleneck in the first ~50-ms of light. Moreover, the steady state rates were elevated in both strains when nigericin was present. In these cells, P700 redox states were changed upon nigericin addition (Figure 4D). In the absence of the drug, the control strain established a larger YNA and thus a smaller YI than *PETC-P171L*. The YND parameter was lowered to similar levels after adding nigericin. Upon nigericin addition, *PETC-P171L* cells established YNA and YI like control. Moreover, samples were indistinguishable in the absence of PSII activity (HA+DCMU, Figure 4D).

Taken together, these spectroscopic findings reveal that the P700 redox states under the actinic light illumination are differently distributed in *PETC-P171L* cells when oxygen levels were low. The different P700 redox state in *PETC-P171L* relied on upstream events on the level of the cytochrome *b*_6_*f* and, as shown in single-turnover measurements, the ΔpH component of the *pmf* could be identified as being one of the limiting factors in oxygen-deprived algae. In most experiments at ~160 μmol photons·m^−2^·s^−1^ actinic red light, *PETC-P171L* performed like control strains when cells were well aerated, suggesting that the acidification of the lumen pH in the permissive light was insufficient to trigger the cytochrome *b*_6_*f* phenotype. In the last two results sections, we will explore the *PETC-P171L* performance in the presence of oxygen at various light intensities.

### The *PETC-P171L* mutant showed an impaired photoautotrophic growth rate under high light

We evaluated whether the *PETC-P171L* mutation has an impact on cellular growth over the course of days. The control and *PETC-P171L* cells grown in liquid medium were spotted on solid medium and the cell growth was observed (Figure 5A). The cells showed a similar photoheterotrophic growth under moderate light illumination on solid TAP medium. On the other hand, when cells were grown photoautotrophically on solid TP medium, *PETC-P171L* cells grew more slowly than control cells under illumination of 100 μmol photons·m^−2^·s^−1^ and 600 μmol photons·m^−2^·s^−1^. Although the *PETC-P171L* cells were able to grow photoautotrophically, the relative growth rate of the mutant cells was apparently slower in the stronger light (Figure 5A). We also analyzed the photoautotrophic growth of control and *PETC-P171L* strains in liquid medium under illumination of 200 μmol photons·m^−2^·s^−1^. We monitored the chlorophyll concentration of liquid culture as a measure of cell growth (Figure 5B). The doubling times of control and *PETC-P171L* were estimated to be 9.4 and 14 hours, respectively, indicating that *PETC-P171L* cells grow 1.5 times slower under 200 μmol photons·m^−2^· s^−1^.

**Figure 5.**
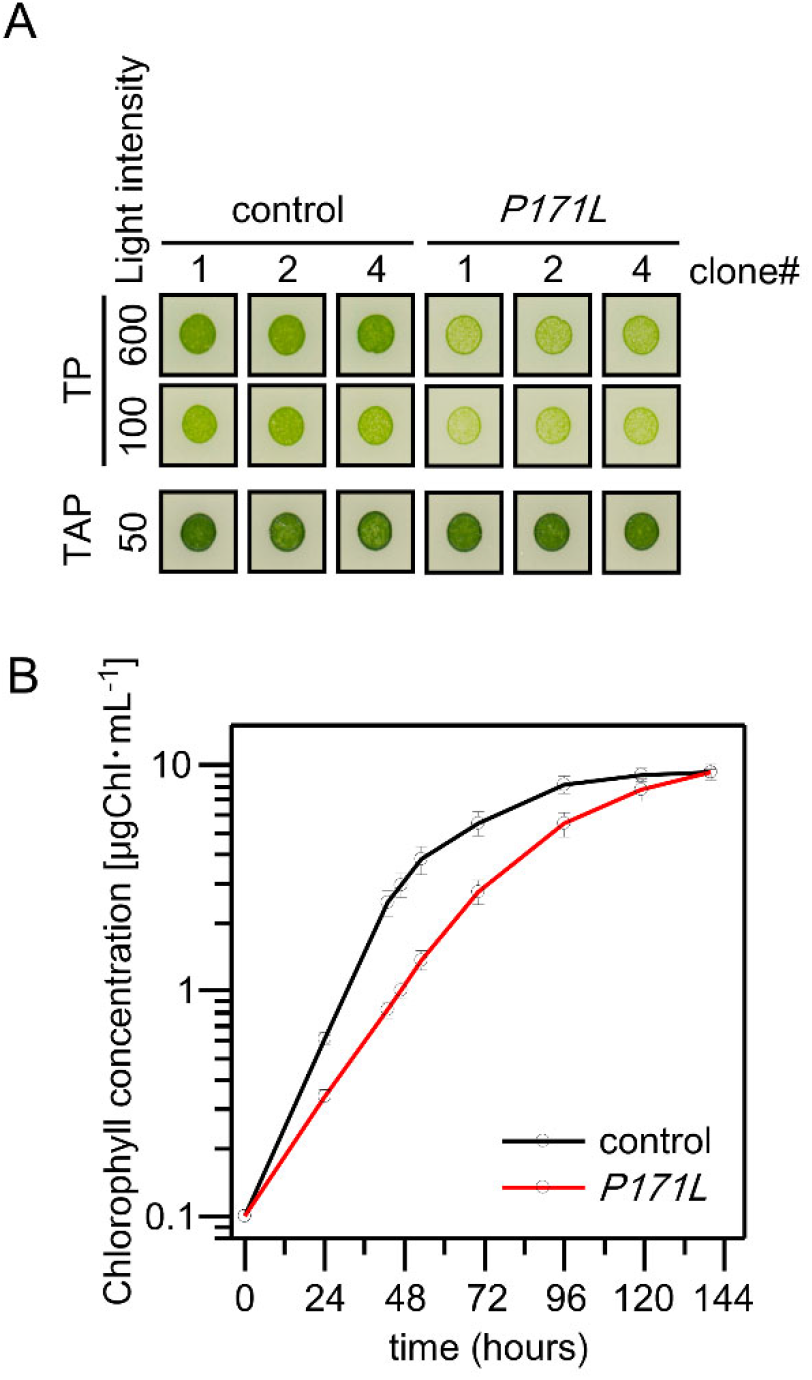
The photoautotrophic growth rate is slowed down at low cell density in *PETC-P171L*. **(A)** Cells were grown photoheterotrophically on solid TAP medium (50 μmol photons·m^−2^·s^−1^) or photoautotrophically on solid TP medium (100 or 600 μmol photons·m^−2^·s^−1^). Three independent clones of the control and *PETC-P171L* strain (*P171L*) were analyzed. **(B)** Photoautotrophic cell growth is monitored by measuring Chlorophyll concentration of the TP liquid culture medium grown under continuous light (200 μmol photons·m^−2^·s^−1^). Black line represents control and red line represents *PETC-P171L* (*P171L*). Three biological replicates were averaged (±standard errors).

### The *PETC-P171L* shows normal state transitions but displays an impaired NPQ induction under strong light illumination and during CO_2_ limitations

The slower photoautotrophic growth of *PETC-P171L* cells may be ascribed to higher photosensitivity at an intensity of 200 μmol photons·m^−2^·s^−1^. One photoprotection strategy involves NPQ to which state transitions are significantly contributing to in algae (Allorent et al., 2013). A functional Qo-site is required to activate the STT7 kinase for catalyzing state transitions (Vener et al., 1997). Thus, the higher photosensitivity in *PETC-P171L* cells may result from the point mutation disturbing state transitions. Therefore, we measured fluorescence emission spectra at 77 K after the induction of state transitions. Indeed, control and the *PETC-P171L* cells exhibited higher intensity of PSII core fluorescence emission (peaking at 688 nm) in state 1 while the PSI core fluorescence emission (peaking at 716 nm) was higher in state 2 (Figure 6A). This indicates that both state 1 and state 2 were fully induced and that the *PETC-P171L* mutation does not affect state transitions. Moreover, we tested whether the accumulation of the LHCSR3 protein is affected since it is necessary to establish the energy-dependent NPQ in algae (Peers et al., 2009). We observed that LHCSR3 was similarly induced after 4 h of strong light illumination in control and *PETC-P171L* cells grown photoautotrophically (Figure 6B). Accordingly, the requirements to establish NPQ are met in *PETC-P171L*.

**Figure 6.**
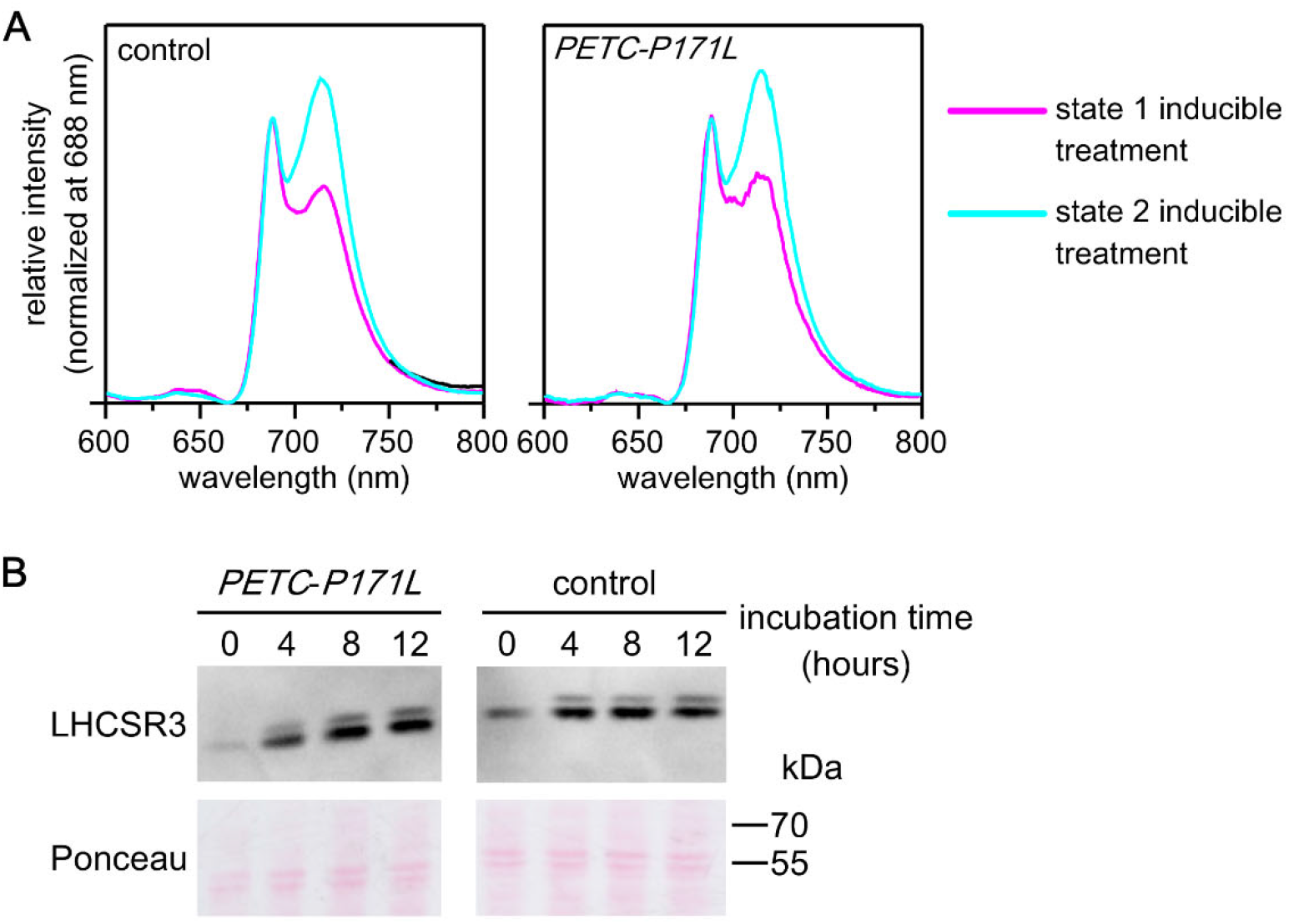
The *PETC-P171L* mutation does neither affect induction of state-transitions, nor LHCSR3 expression induction and/or stability. **(A)** The TP-grown Chlamydomonas cells acclimated under high light (300 μmol photons·m^−2^·s^−1^) were resuspended with TP at 2 μg Chlorophyll·mL^−1^ and incubated in state 1 or state 2 inducible condition. In the case of state 1 inducible treatment, cells were incubated under 200 μmol photons·m^−2^·s^−1^ in the presence of DCMU for 10 minutes. In the case of state 2 inducible treatment, cells were incubated in the darkness for 10 minutes in the presence of 100 mM glucose and 50 U·mL^−1^ glucose oxidase. Samples were excited with 430 nm and the emitted fluorescence were recorded from 600 to 800 nm in liquid nitrogen. The fluorescence spectra were normalized at 688 nm. **(B)** The cells grown in TP medium were shifted from low light (15 μmol photons·m^−2^·s^−1^) to stronger light (200 μmol photons·m^−2^·s^−1^) and immune-chase experiment with the antibody against LHCSR3 protein was performed for cellular proteins. Cells were harvested before shifting to higher light (0), after 4, 8, and 12 hours shifting to high light. Ponceau staining panel served as loading control.

Next, we monitored photosynthesis via chlorophyll fluorescence over a 16-h photoperiod to seek for a link between a possible restriction in the *PETC-P171L* photosynthetic machinery and the growth phenotype that we observed in Figure 5. Therefore, we cultivated the control strain and *PETC-P171L* in photobioreactors in the absence of air supply to favor CO_2_ limitation and induce NPQ (Kanazawa and Kramer, 2002). We followed the effective PSII quantum yield (Y(II)) and NPQ for 72 h (Supplemental Figure 9), and the progressive Y(II) decline at dawn was more pronounced in *PETC-P171L*. Figure 7 displays a typical 16-h photoperiod peaking at 250 μmol photons·m^−2^·s^−1^. The low Y(II) of ~0.1 during elevated light levels was similar in both strains (Figure 7A). However, the initial drop in Y(II) was more pronounced in *PETC-P171L* during the first 2 h of light, and the relaxation from lower to higher Y(II) values at the end of the photoperiod was slightly delayed in *PETC-P171L*, reaching stable Y(II) after ~1 h darkness. The associated NPQ values differed more significantly beyond the first 2 h of the photoperiod (Figure 7B). They were strongly reduced to ~0.4 in *PETC-P171L*. NPQ peaked at 1.1 in the control strain after 4 h of light, slowly decreasing to 0.8 at the end of the light period. Moreover, NPQ relaxation after the end of the light period (after 41 h of culture time in the figure) was slower in *PETC-P171L* than in the controls (~2 h vs 1 h). Our photobioreactor findings are quantitatively in agreement with additional PAM fluorimetry measurements. Therein (Supplemental Figure 10), we observed that high light treated *PETC-P171L* cells showed a lower Fv/Fm and, especially in low light, a smaller Y(II) due to a compromised NPQ formation efficiency that paralleled a more reduced PQ pool.

**Figure 7.**
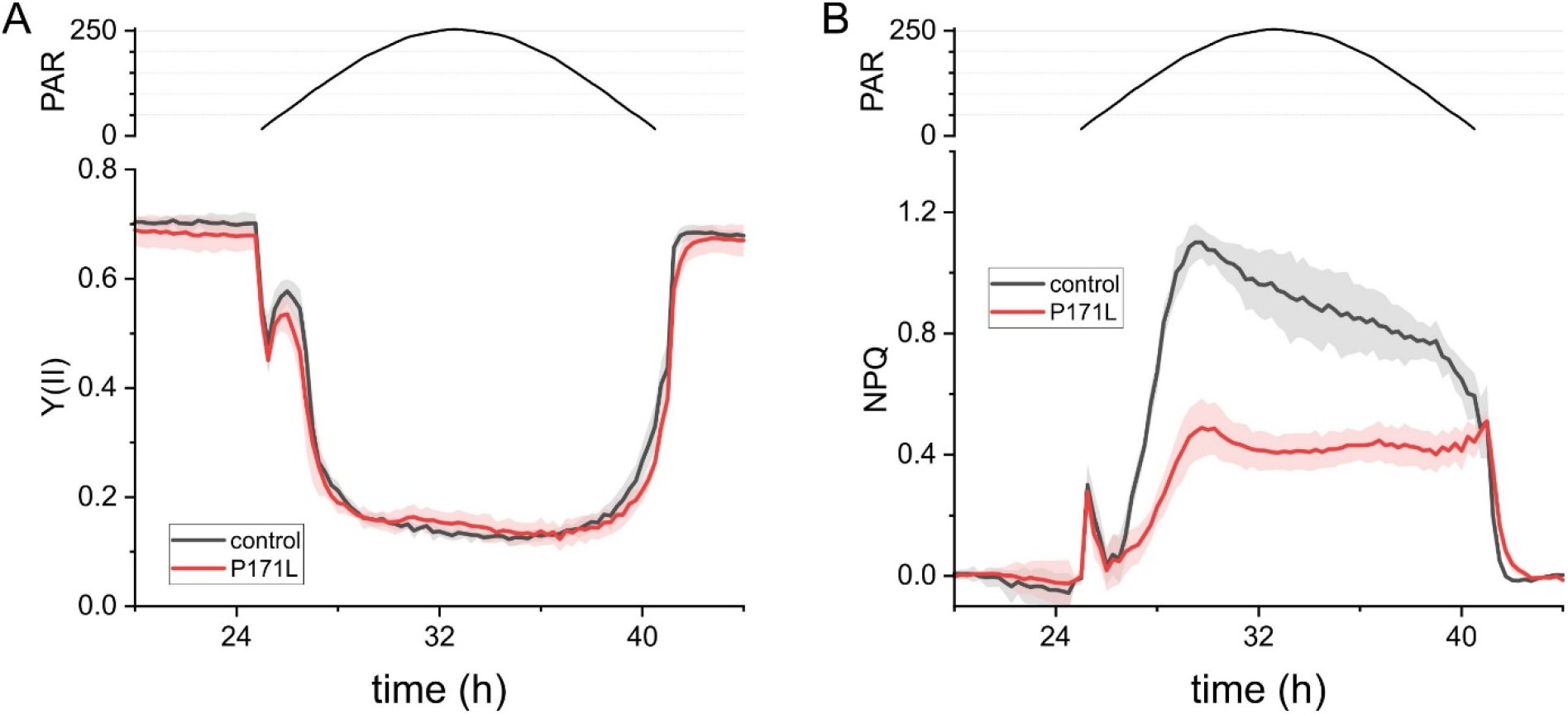
CO_2_ limitation reveals the chlorophyll fluorescence phenotype of a limited photosynthetic capacity in *PETC-P171L* at lower irradiances. Averaged photobioreactor experiments (n = 3 ±SD) of non-aerated cultures are shown for control and *PETC-P171L* cells. **(A)** The PSII quantum yield Y(II) and **(B)**NPQ show that *PETC-P171L* performance is less efficient in the twilight and throughout the day, respectively. The second photoperiod is shown (16-h light / 8-h darkness, PAR is μmol photons·m^−2^·s^−1^, for complete experiments refer to Supplemental Figure 9).

Taken together, the chlorophyll fluorescence parameters in *PETC-P171L* suggested a limitation of photosynthesis on the level of cytochrome *b*_6_*f* complex at elevated irradiances and/or when CO_2_ availability is low.

## Discussion

In Arabidposis, the *pgr1* mutant showed reduced NPQ owing to cessation of total electron transfer at elevated light intensities above ~200 μmol photons·m^−2^·s^−1^, most likely at the level of *b*_6_*f* (Munekage et al., 2001; Okegawa et al., 2005). P700 redox measurements of isolated *pgr1* thylakoids further suggested that the pH-dependent inactivation of electron transfer via the *b*_6_*f* was shifted towards more alkaline pH by about 1 unit (Jahns et al., 2002). Yet, direct *b*_6_*f* measurements of the Arabidopsis *pgr1* mutant are missing in literature. In the current work we successfully generated a *PETC-P171L* strain in *C. reinhardtii*, harboring a genetic site-directed alteration corresponding to the *pgr1* mutation in *A. thaliana* (Shikanai et al., 1999). In *C. reinhardtii*, this mutation did neither affect *b*_6_*f* accumulation, state transitions, nor genetic NPQ regulation via LHCSR induction. We directly show that, depending on the environmental conditions, a ΔpH-dependent slow-down of electron transfer through the algal *PETC-P171L b*_6_*f* complex is established. This restriction most likely explains the observed growth defect in the mutants due to diminished electron transfer activity. In the algal system, under conditions where CO_2_ limitation was initiated in sealed photobioreactors, the algal Rieske mutant showed a lower NPQ above ~100 μmol photons·m^−2^·s^−1^ already.

### The PETC-P171L mutation slows down the electron transfer through *b*_6_*f* in lumenal pH acidification

The previous studies observed strong NPQ reduction in the Arabidopsis *pgr1* mutant in high light. This phenotype linked to reduction of electron transfer in the mutant plants (Munekage et al., 2001; Okegawa et al., 2005; Yamamoto and Shikanai, 2019). Our measurements of photosynthetic control in *PETC-P171L* revealed similarities to the available data in vascular plants. Namely, the bottleneck of electron transfer in *PETC-P171L* was detected at the level of PSII via chlorophyll fluorescence (Figure 7) and, depending on the condition, at the level PSI (Figure 3B). Moreover, we ruled out that state transitions were affected by the Qo-site mutation (Figure 6A) and further evaluation of the photosynthetic machinery pinpointed the *PETC-P171L* defect to the *b*_6_*f* complex (Figure 4). We identified a significant *b*_6_*f* defect in *PETC-P171L* under anoxia which had an impact on cytochrome-*f* reduction, Q-cycle electrogenicity, faster electron flow cessation during photosynthetic induction and facilitated P700 oxidation in the light. This relates the *PETC-P171L* phenotype for the first time to a mechanistic defect in the *b*_6_*f* complex that is due to an altered pH-dependence within the mutant *b*_6_*f* complex. Accordingly, the addition of nigericin, exchanging ΔpH for ΔΨ, abolished the impact of the mutation. Anaerobiosis increases the ΔpH contribution to the *pmf* in darkness, which depends on functional ATP synthase (Finazzi and Rappaport, 1998; Rappaport et al., 1999). Oxygen deprivation caused a charge separation deficiency of the *b*_6_*f* in *PETC-P171L*, which is a consequence of delayed mutant Qo-site turnover. However, our nigericin approach to address the ΔpH impact could not restore the same rapid charge separation onset in *PETC-P171L* that we observed in the control strain (Figure 4B). Thus, the mutation could affect the *b*_6_*f* complex on multiple levels. In fact, a cooperativity of high- and low-potential chain reactions has been documented in previous Rieske iron sulfur protein (ISP) mutants of the cytochrome *bc*_1_ complex from yeast (Snyder et al., 2000). Still, this coupling mechanism operating between Qo- and Qi-site remains elusive. The authors mutated homologous Rieske ISP residues at position 157 and 159 (orange-colored rectangle in Supplemental Figure 3), not too far from proline 171 of *C. reinhardtii* PETC, which eventually result in a more reducing redox midpoint potential (*E*_m_) of the Rieske ISC and a slowdown of *b*-heme reduction (Snyder et al., 2000). Thus, similar effects could be anticipated in *PETC-P171L* provided that the mutation lowers the Rieske ISC *E*_m_ in *b*_6_*f* as originally proposed (Munekage et al., 2001; Jahns et al., 2002). However, the *E*_m_ has not been determined in *pgr1*. Previous data indicated that the *E*_m_ of reconstituted recombinant *C. reinhardtii* Rieske ISC is pH-dependent with a pK_ox_ of 6.2 and a pK_red_ of 8.2 (Soriano et al., 2002). The *E*_m_ changes in the pH-dependent region with −40 mV/pH. Thus, lowering the Rieske ISC *E*_m_ via the PETC-P171L mutation could impact pK_ox_ and pK_red_, and might alter inter *b*_6_*f* electron transfer as shown in yeast (Snyder et al., 2000). In an alternative scenario, the PETC-P171L mutation might alter binding of PQH_2_ to the Q_o_-site and consecutively change the pH-dependent redox properties and oxidation of PQH_2_. The Rieske ISC is found in the hydrophilic domain of PETC (Figure 1B) and, based on structural similarities with *bc*_1_ complex (Darrouzet et al., 2000), it is supposed to be a flexible hydrophilic domain to realize electron transfer between redox centers Rieske ISC and heme *f* Additionally, proline 171 mutated in this study is the second proline of the “Proline-X-Proline” motif, which is a common structure at the beginning of turn structure (MacArthur and Thornton, 1991). This motif is exclusively found in PETC protein and conserved among Rieske ISP (Supplemental Figure 3). Accordingly, this motif forms a typical hairpin structure and the substitution of the second proline to leucine may disturb conformation of PETC hydrophilic domain structure, causing the functional impact in the redox chemistry of *b*_6_*f*.

Our findings revealed that the generated ΔpH in anaerobic Chlamydomonas during 30-s of darkness was sufficient to trigger severe photosynthetic control in *PETC-P171L*, preventing efficient electrogenicity via the low-potential chain due to aggravated cytochrome-*f* reduction. As the effect of nigericin on photosynthetic control are well established (Joliot and Johnson, 2011), our nigericin data also revealed a significant contribution of the ΔpH in anaerobic cells (Figures 3 and 4). Accordingly, *b*_6_*f* regulation is important for PSI photoprotection as well as for qE quenching in a reducing cellular environment such as anaerobiosis. The differential regulation of pH dependent inactivation of electron transfer via the *b*_6_*f* in oxic and anoxic conditions in *PETC-P171L* is surprising. This might suggest that depending on oxygen availability (or metabolic congestion), the regulation of photosynthetic control is distinct in *C. reinhardtii*. However, revealing a low-light *pgr1* phenotype may also occur in plants when metabolism becomes restricted, e.g. under drought stress conditions where a downregulated ATP synthase activity favors large ΔpH amplitudes (Kohzuma et al., 2009). How can this be rationalized? Three non-exclusive scenarios may occur: First, metabolic control of ATP synthase downregulates H^+^ conductivity which theoretically facilitates a light-dependent ΔpH build up in anaerobic conditions. Second, sustaining a ΔpH in an ATP synthase-dependent manner is favored in anoxic algae even in darkness (Finazzi and Rappaport, 1998; Rappaport et al., 1999). Third, the regulation of *b*_6_*f* electron transfer via the Q-cycle is distinct under oxic and anoxic conditions (Buchert et al., 2020; Buchert et al., 2022). Under anoxic conditions stromal electron intake into *b*_6_*f* changes the Q-cycle from a canonical cycle during LEF to an alternative cycle during CEF, which would facilitate more efficient proton translocating into the lumen as compared with the canonical Q-cycle. In line, photosynthetic control would be faster activated under anoxia and revealed the *PETC-P171L* phenotype even in low-light conditions.

Taken together, *PETC-P171L* may be a useful tool for future investigations under various metabolic and environmental conditions to uncover details of the cooperative mechanism between Qo- and Qi-site. Moreover, *PETC-P171L* may help to elucidate the interplay of various *pmf*-shaping ion channels/antiporters involved to fine-tune *pmf* partitioning in algae in response to metabolic constraints.

### PETC-P171L mutation retards photoautotrophic cell growth

The growth phenotype of *pgr1* plants under different light intensities has not been addressed in literature (Shikanai et al., 1999; Munekage et al., 2001; Yamamoto and Shikanai, 2019). We found that *PETC-P171L* grew less efficiently under photoautotrophic condition at growth rates 1.5 times slower in the logarithmic phase than control (Figure 5 and Supplemental Figure 11). Despite similar amounts of LHCSR3 in the control and *PETC-P171L* cells, NPQ could not be fully established in *PETC-P171L* and PSII performance was low due to a more reduced PQ pool in the mutant (Figure 7, Supplemental Figure 10). This analysis shows the *PETC-P171L* mutation impacts cellular growth and underlines the importance of a balanced photosynthetic control in wild type cells (control strain), including qE regulation via acidification of the lumen. These data demonstrate the tight interconnection of pH-dependent regulation of photosynthetic control and qE driven quenching, which also requires expression of LHCSR3.

Taken together, *PETC-P171L* exhibits a reduced photosynthetic electron transfer rate because of its hypersensitive photosynthetic control mechanism. The light intensity threshold to trigger the hypersensitive effect in the mutant *b*_6_*f* can be modulated by the metabolic state of the cell. The molecular machinery to establish photosynthetic control, energy quenching and balancing of light harvesting requires functional wild type *b*_6_*f* strains. These results are consistent with Arabidopsis studies, indicating that the effect of the proline substitution in PETC on electron transfer is *per se* conserved between the two organisms. Moreover, our study proves that the *PETC-P171L* phenotype is directly linked to functional properties of the *b*_6_*f* complex.

## Acknowledgement

The *petC-Δ1*strain and pACR4.5 plasmid were kindly gifted by Catherine de Vitry. The cDNA library of *C. reinhardtii* and the pSL18 plasmid expression vector were provided by Michel Goldschmidt-Clermont. The antibodies against PsaA and PetA/PETC were kindly gifted by Kevin Redding and Olivier Vallon, respectively.

